## Supplemental figures for "OsNF-YB7 inactivates OsGLK1 to inhibit chlorophyll biosynthesis in rice embryo"

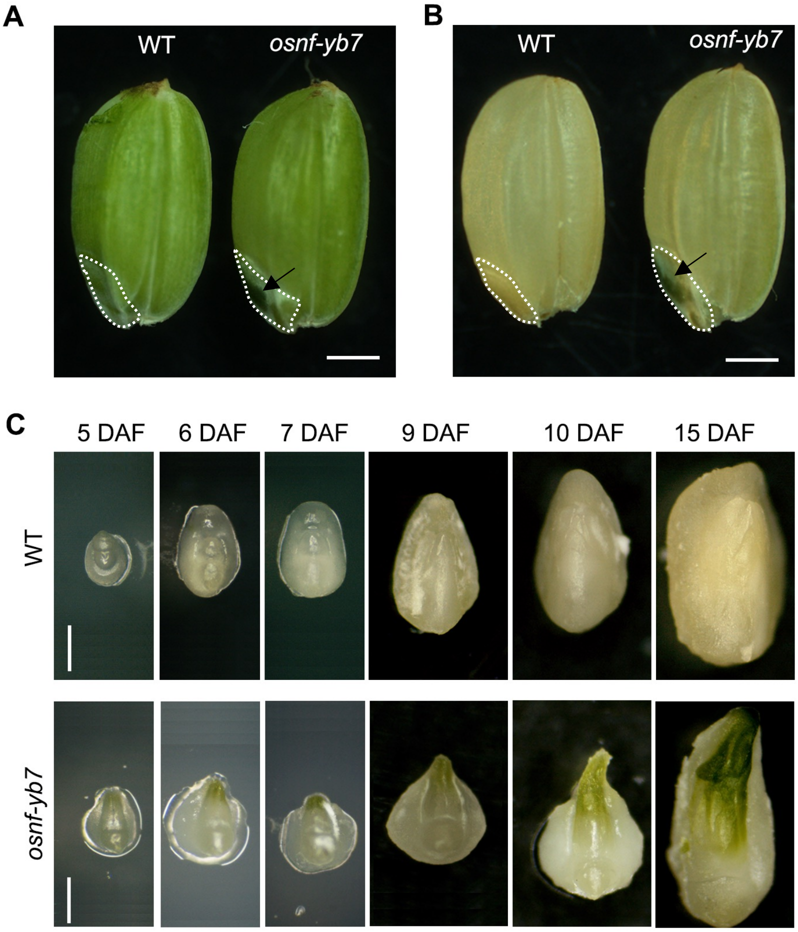


**Supplementary Figure 1. Mutation of *OsNF-YB7* leads to chloroembryo.**
**A, B.** Morphologies of WT and *osnf-yb7* caryopses at 20 DAF (**A**) and at maturation (**B**). The embryos are circled with dash lines, arrow heads indicate green embryos produced by *osnf-yb7*. Bars = 1 mm.
**C.** Images showing the embryo development of WT and *osnf-yb7* at different stages. Bars = 1 mm.

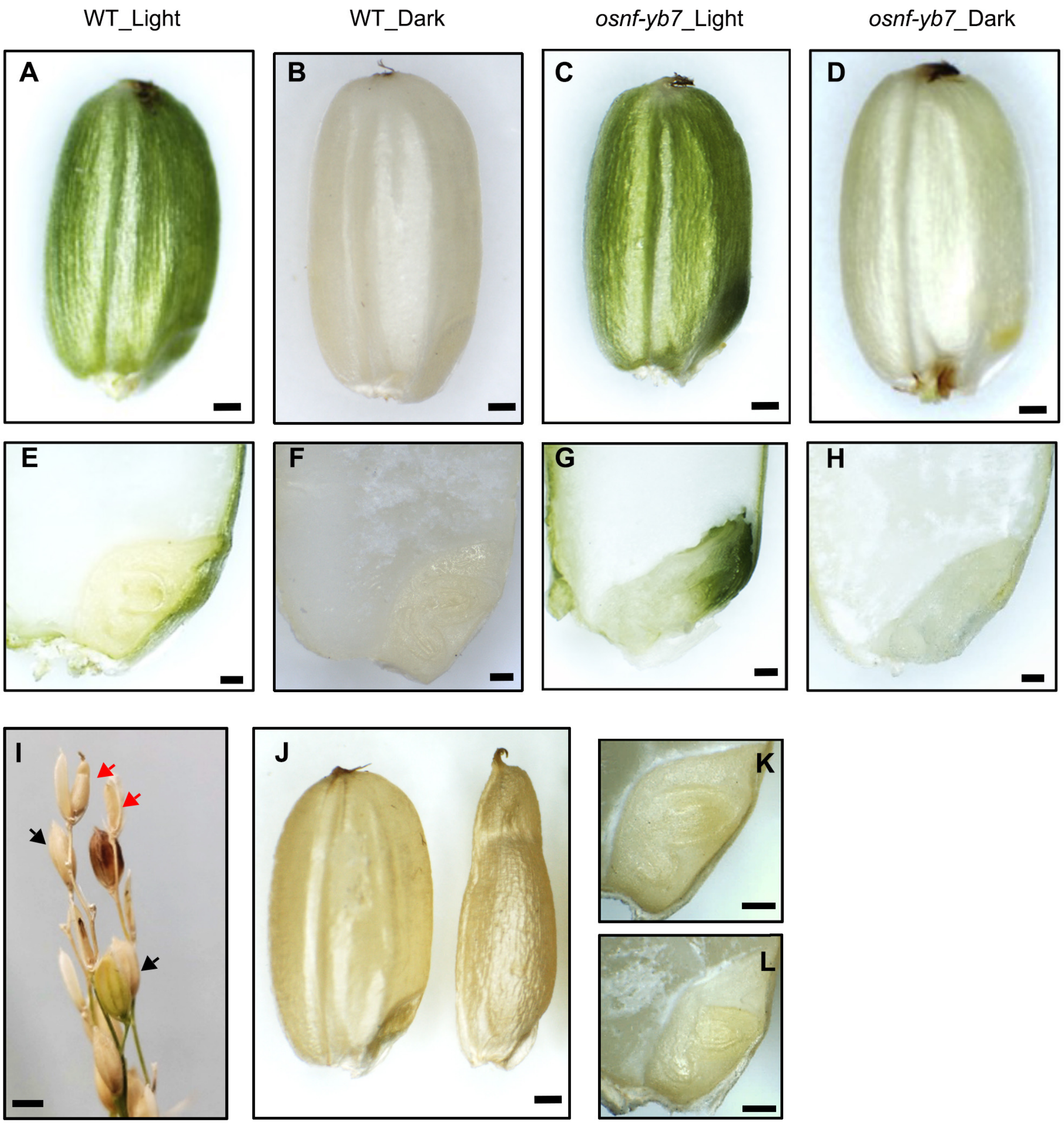


**Supplementary Figure 2. Light is required but not sufficient for Chl biosynthesis in the chloroembryo of *osnf-yb7*.**

**A–D.** Phenotype of the WT (**A, B**) and *osnf-yb7* (**C, D**) caryopses that grew under normal (**A, C**) and dark-treated (**B, D**) conditions. Scale bars = 500 μm.

**E–H.** Phenotype of the WT (**E, F**) *and osnf-yb7* (**G, H**) embryos that grew under normal (**E, G**) and dark-treated (**F, H**) conditions. The embryos were longitudinally dissected. Scale bars = 500 μm.

**I.** An image showing the mature seeds whose hulls were surgically removed soon after fertilization (approximately 1 to 2 DAF) to allow the embryos expose to light. The red and black arrows indicate the seeds with the hulls removed or not removed, respectively. Scale bar = 5 mm.

**J.** Phenotype of WT seed grew under normal condition (left) or with the hulls removed (right). Scale bars = 500 μm.

**K, L.** Longitudinally dissected embryos of WT that grew under normal condition (**K**) or with the hulls removed (**L**). Scale bars = 500 μm.

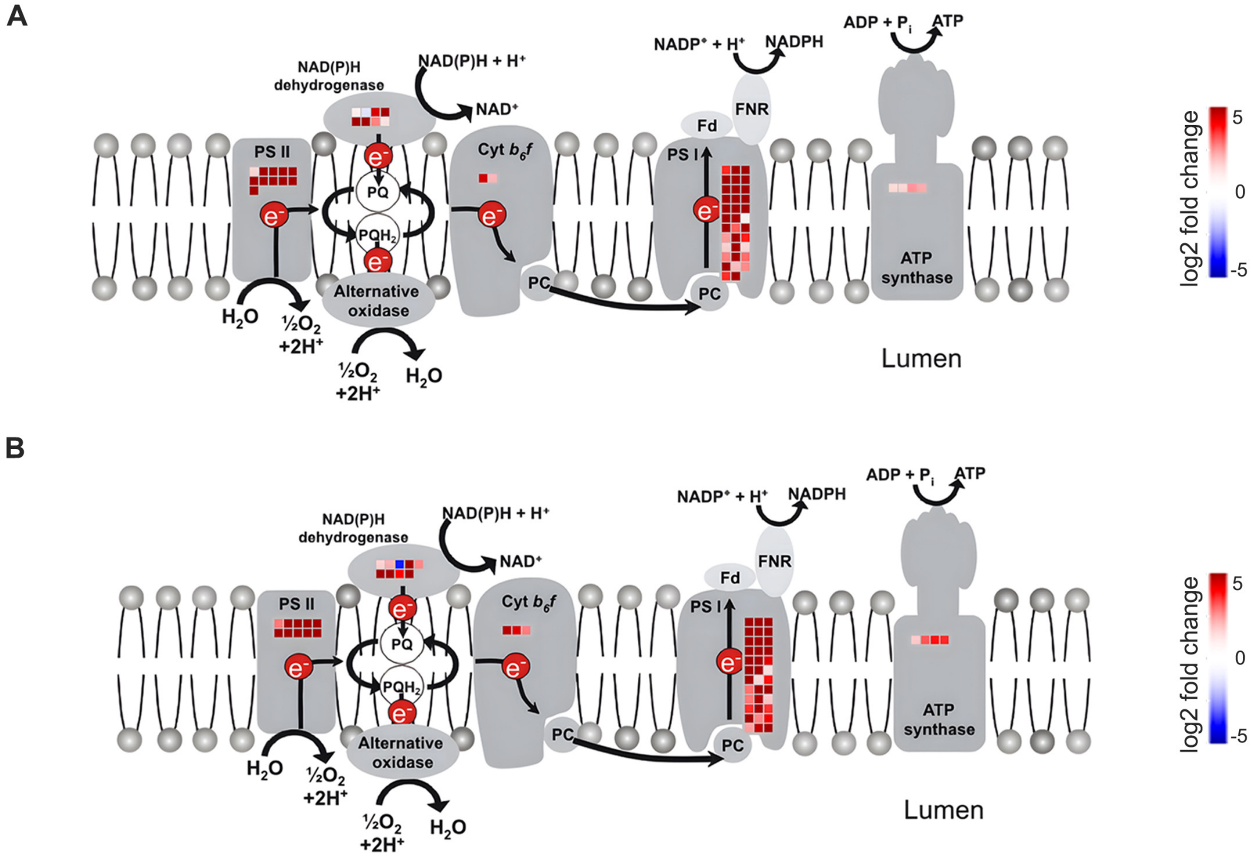


**Supplementary Figure 3. Mutation of *OsNF-YB7* actives the expression of photosynthesis-realted genes.**

**A, B.** Schematic illustration of the activated photosynthesis-related genes in the *osnf-yb7* embryos at 5- (**A**) and 10 DAF (**B**)

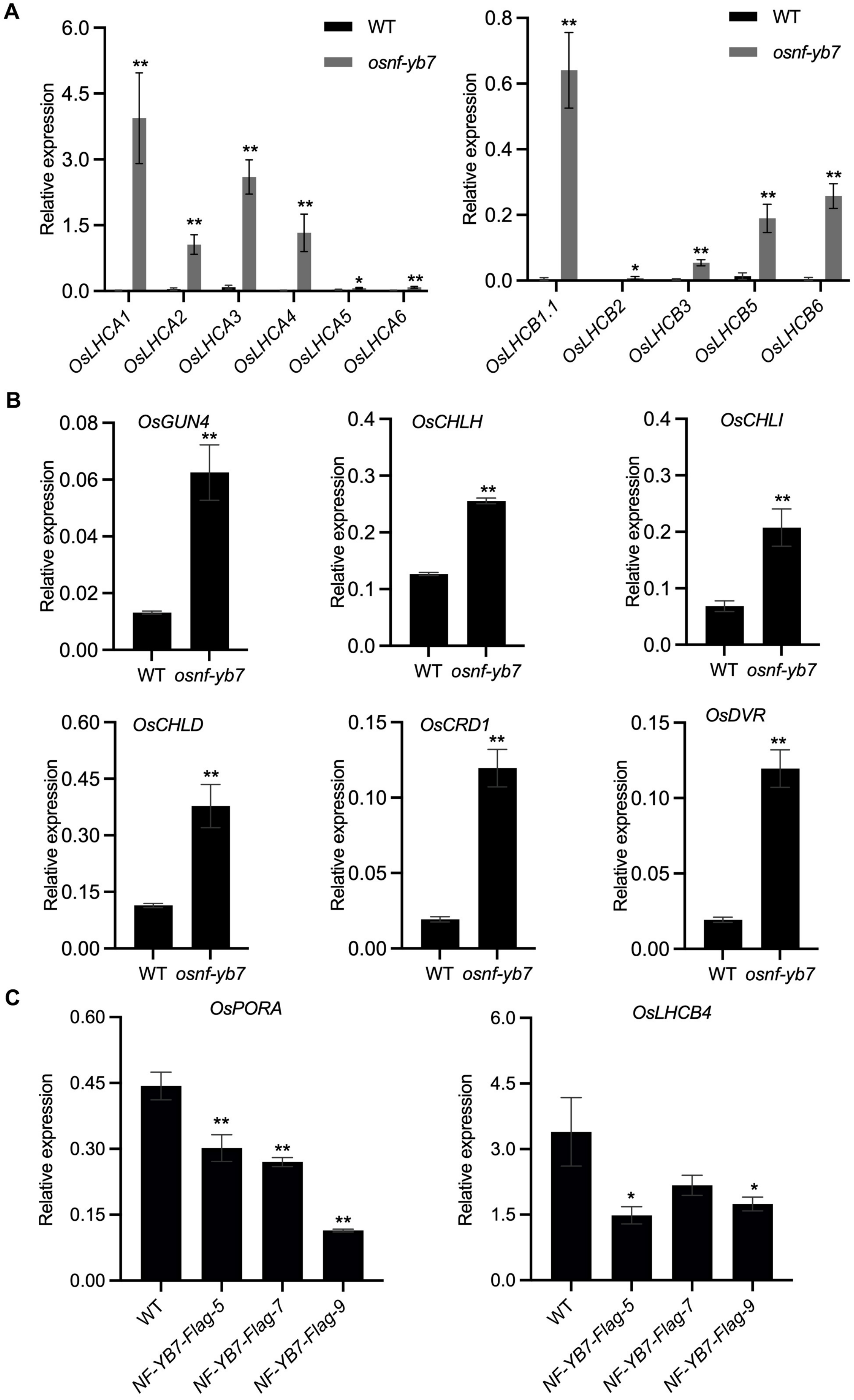

A
WT
osnf-yb7
Relative expression
OsLHCA1
OsLHCA2
OsLHCA3
OsLHCA4
OsLHCA5
OsLHCA6
OsLHCB1.1
OsLHCB2
OsLHCB3
OsLHCB5
OsLHCB6
B
OsGUN4
OsCHLH
OsCHLI
OsCHLD
OsCRD1
OsDVR
C
OsPORA
OsLHCB4
NF-YB7-Flag-5
NF-YB7-Flag-7
NF-YB7-Flag-9

**Supplementary Figure 4. OsNF-YB7 negatively regulates the expression of light harvest and Chl biogenesis-associated genes.**

**A, B.** RT-qPCR analysis the expression levels of OsLHCs (**A**) and chlorophyll biogenesis-associated genes (**B**) in the embryos of WT and *osnf-yb7* at 10 DAF. Data are means ± SD (n = 3). *, $p < 0.05$; **, $p < 0.01$; Student's *t*-test was used for statistical analysis.

**C.** RT-qPCR analysis the expression levels of *OsPORA* and *OsLHCB4* in the leaves of WT and *B7-OX-Flag* lines. Data are means ± SD (n = 3). *, $p < 0.05$; **, $p < 0.01$; Student's *t*-test was used for statistical analysis.

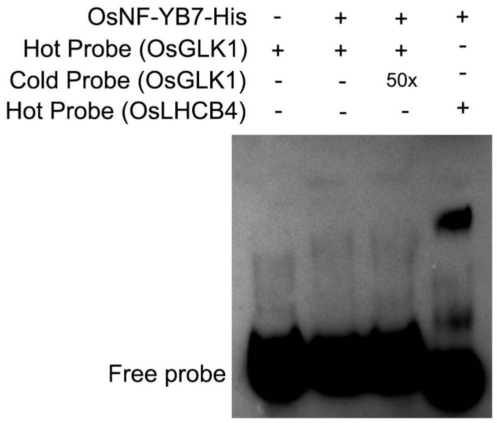


**Supplementary Figure 5. OsNF-YB7 does not directly binds to the promoter of *OsGLK1 in vitro*.**

The EMSA assay showing that OsNF-YB7 did not directly binds to the promoter of *OsGLK1*. OsNF-YB7-His recombinant proteins were able to bind to the promoter of *OsLHCB4*, which was therefore set as a positive control.

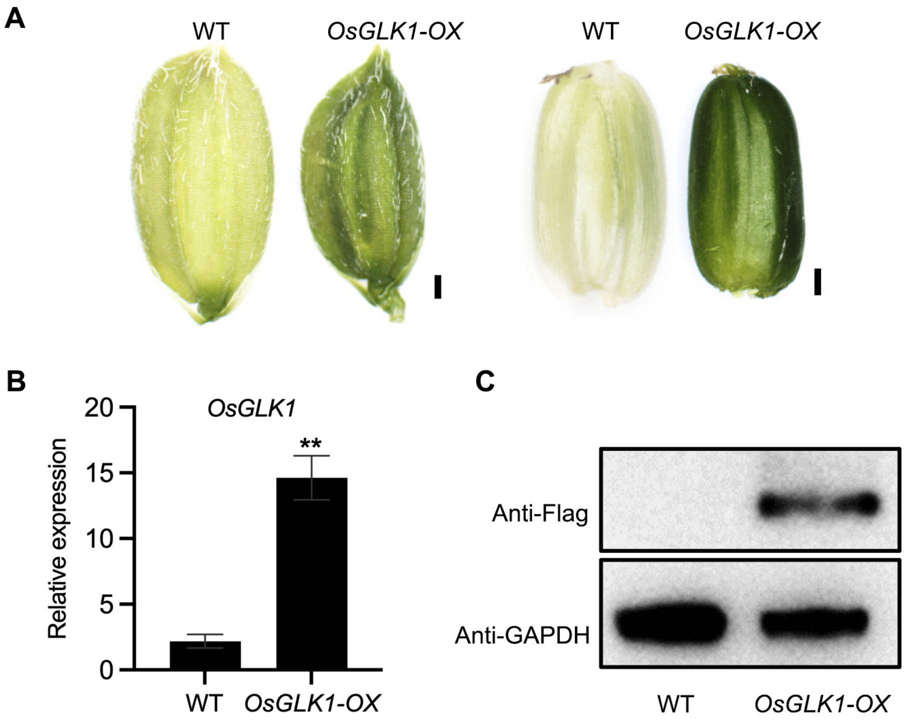


**Supplementary Figure 6. Overexpression of *OsGLK1* induces chloroembryo in rice.**
**A.** Morphologies of the seeds and caryopses produced by WT and *OsGLK1-OX*. Scale bars = 0.5 mm.
**B.** RT-qPCR analysis showing the transcript levels of *OsGLK1* in WT and *OsGLK1-OX*. Data are means ± SD (n=3). **, $p < 0.01$; Student's *t*-test was used for statistical analysis.
**C.** Immunoblot analysis of OsGLK1 proteins in WT and *OsGLK1-OX* using anti-Flag antibody. GAPDH was used to verify that equal amount of proteins were used for loading.

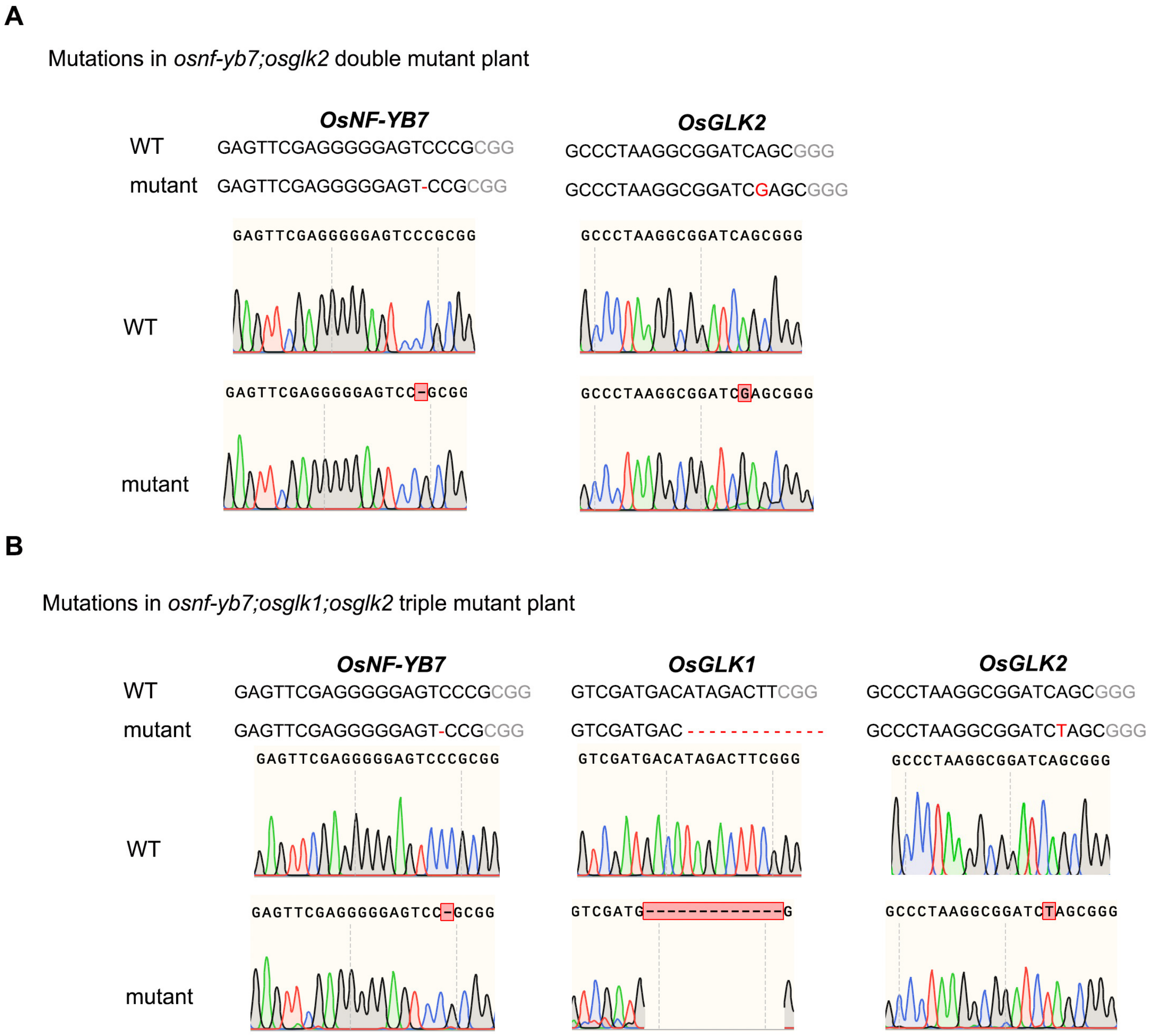


**Supplementary Figure 7. Generation of *osnf-yb7*, *osglk1* and *osglk2* high-order mutants.**
**A.** Sanger sequencing of the *osnf-yb7;osglk2* double mutant.
**B.** Sanger sequencing of the *osnf-yb7;osglk1;osglk2* triple mutant.

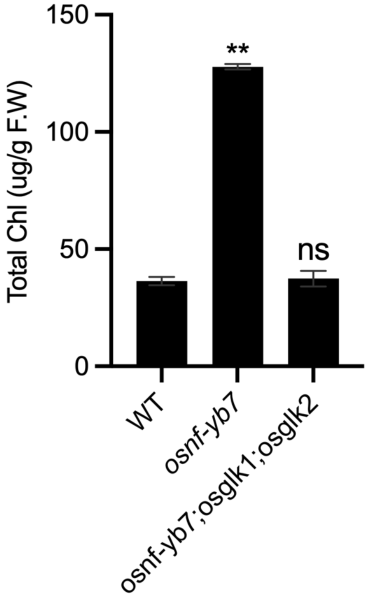


**Supplementary Figure 8. Chl levels in WT, *osnf-yb7* and *osnf-yb7;osglk1;osglk2* embryos at 10 DAF.**
Data are means ± SD (n=3). ns, not significant; **, $p < 0.01$; Student's *t*-test was used for statistical analysis.

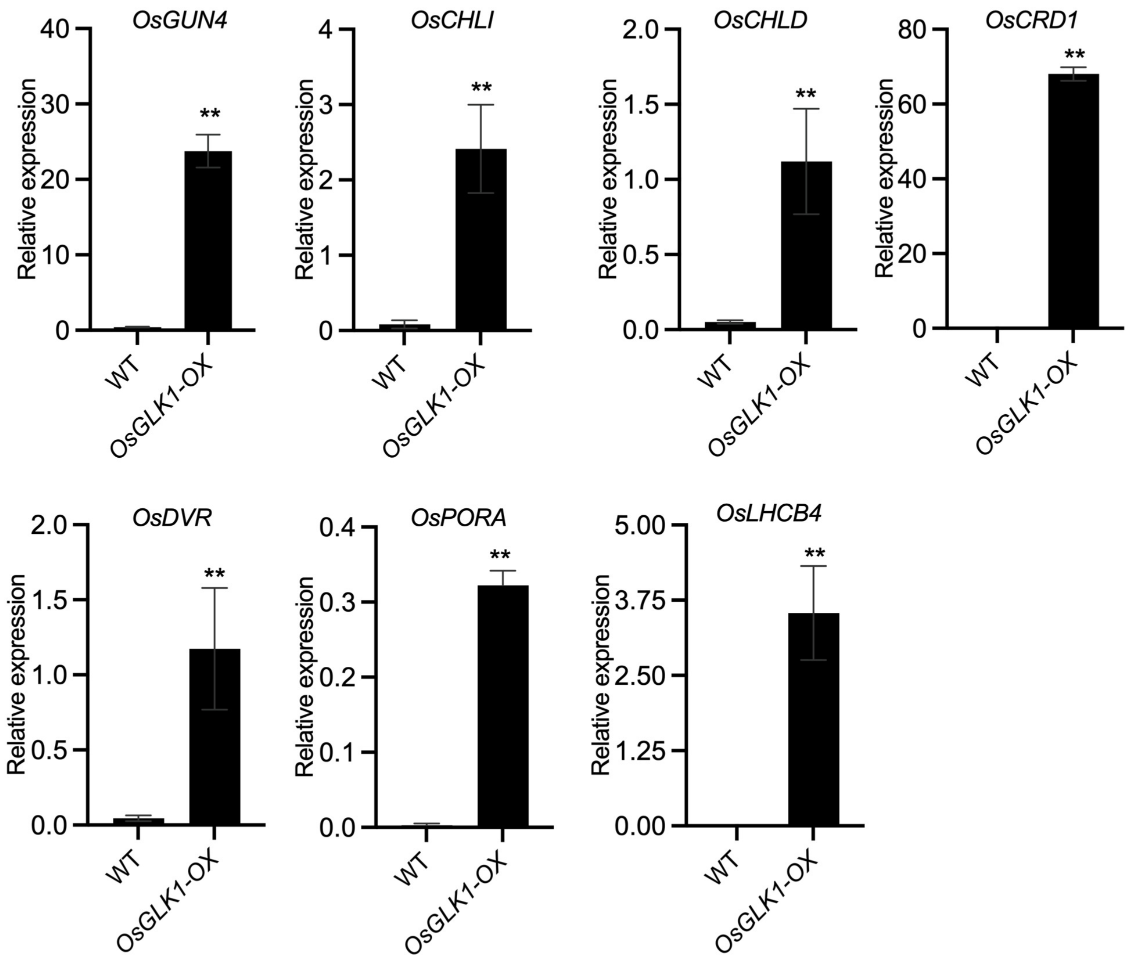


**Supplementary Figure 9. Overexpression of *OsGLK1* activates the expression of Chl biogenesis- and photosynthesis-related genes.**

RT-qPCR analysis of Chl biogenesis- and photosynthesis-related genes. 14-d-old seedlings of WT and *OsGLK1-OX* were used for analysis. Data are means ± SD (n=3). **, $p$ < 0.01; Student's $t$-test was used for statistical analysis.

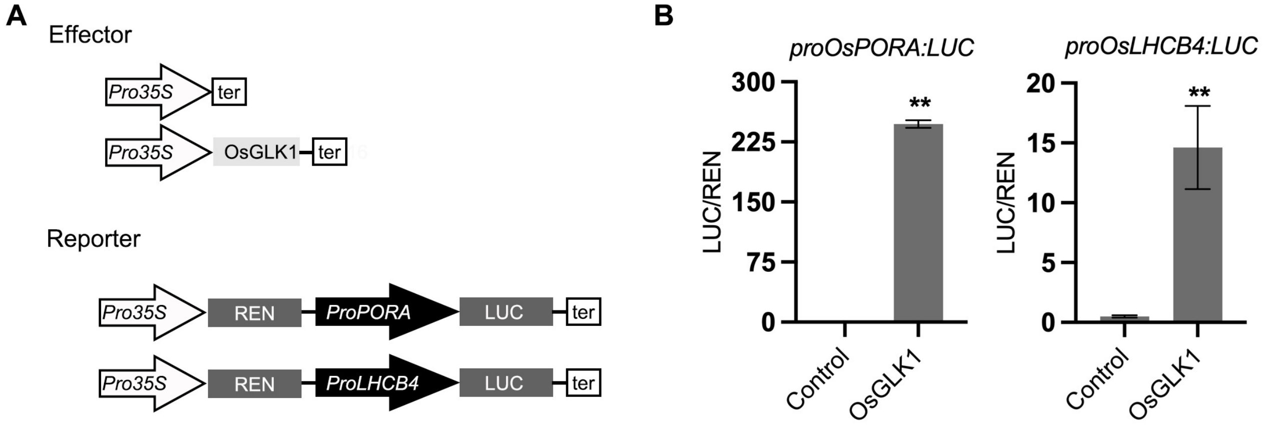


**Supplementary Figure 10. OsGLK1 significantly activates the promoter activities of *OsPORA* and *OsLHCB4*.**

**A.** Schematic diagrams showing the constructs used in the DLR assays. LUC, firefly luciferase; REN, *Renilla* luciferase.

**B.** DLR assays showing that OsGLK1 significantly activated the promoter activities of *OsPORA* and *OsLHCB4* in green tissue protoplasts. **, $p$ < 0.01; Student's $t$-test was used for statistical analysis.

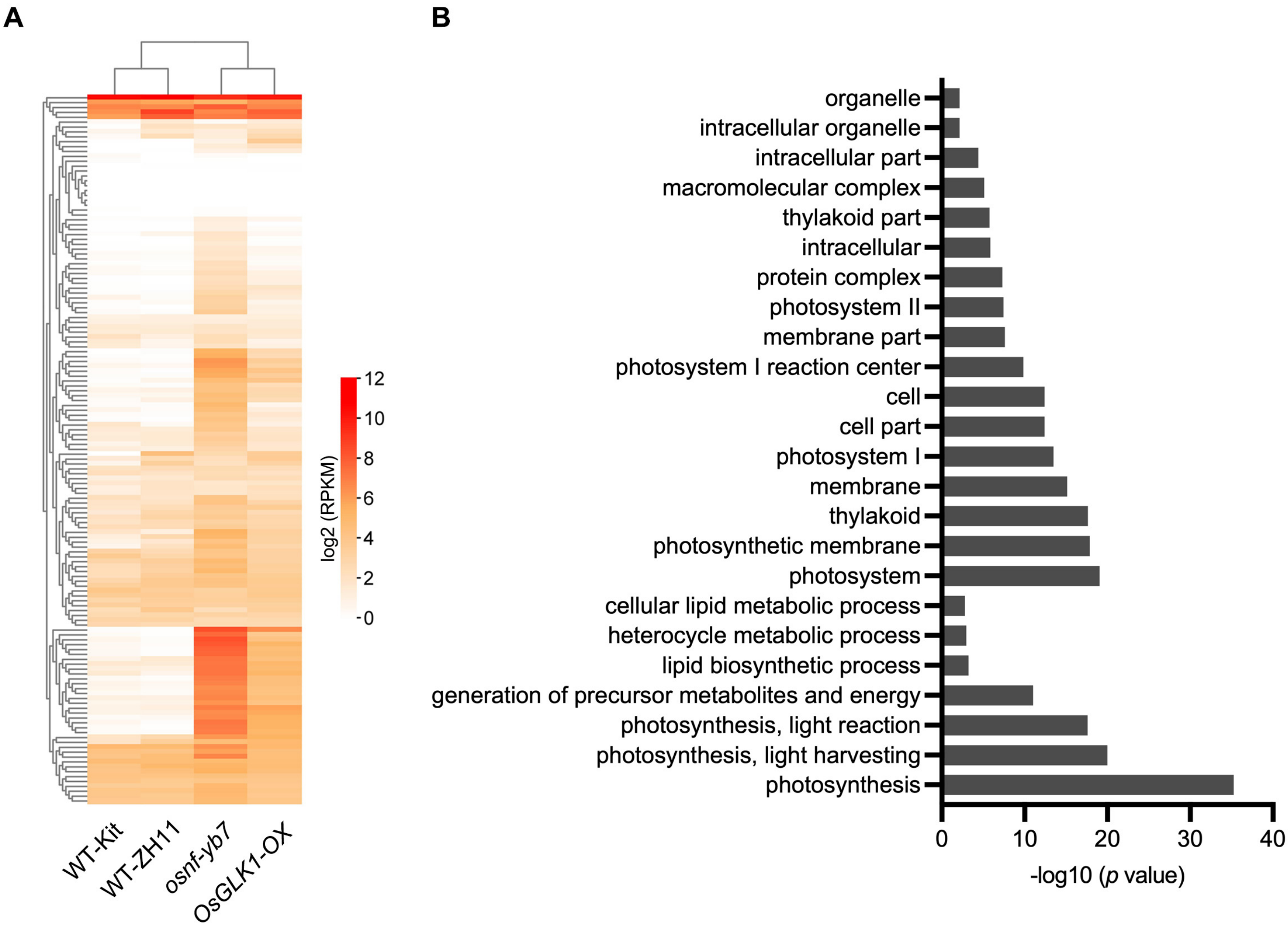


**Supplementary Figure 11. Many of the OsNF-YB7 and OsGLK1 common targets that activated in *osnf-yb7* and *OsGLK1-OX* are involved in Chl biosynthesis and photosynthesis.**

**A.** A heat map showing the expression of the common targets in 10-DAF-old embryos of *osnf-yb7* (Kit background) and *OsGLK1-OX* (ZH11 background) and their corresponding WT.

**B.** Gene Ontology analysis of the common targets of OsGLK1 and OsNF-YB7.

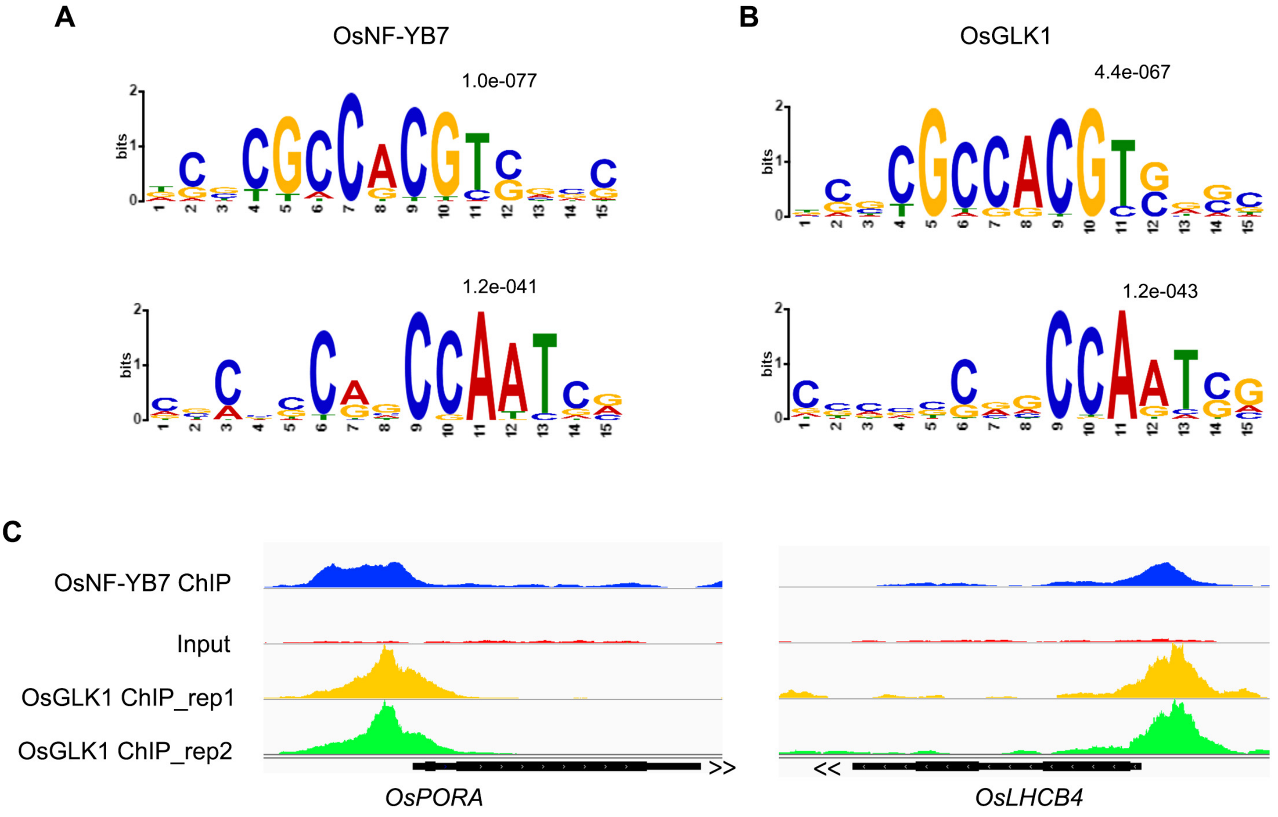


**Supplementary Figure 12. OsNF-YB7 and OsGLK1 bind to proximal regions of their common targets.**

**A, B.** MEME analysis of the motifs that enriched in the binding peaks of OsNF-YB7 (**A**) and OsGLK1 (**B**).

**C.** IGV screenshots showing that OsNF-YB7 and OsGLK1 bind to the same regions of *OsPORA* and *OsLHCB4*.

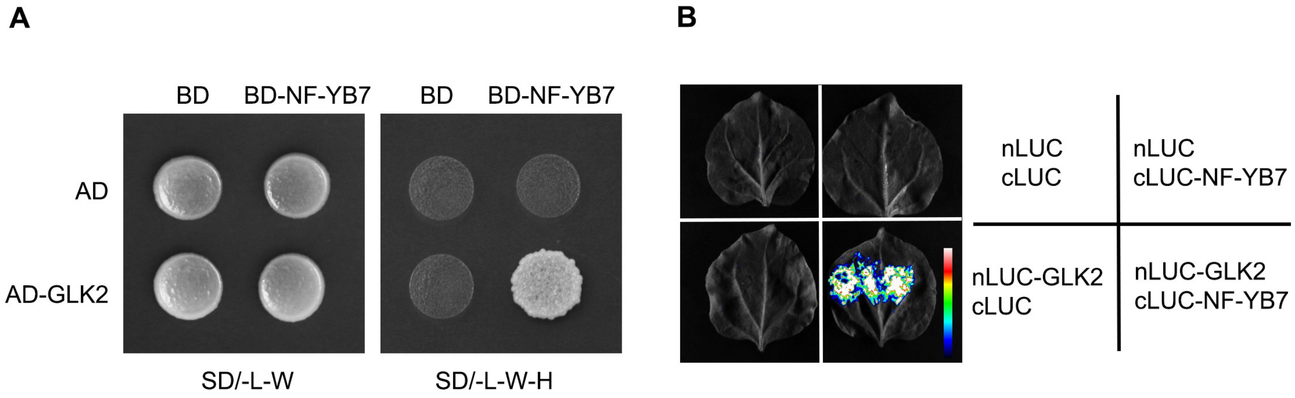


**Supplementary Figure 13. OsNF-YB7 interacts with OsGLK2.**

**A.** OsNF-YB7 interacts with OsGLK2 in yeast. The indicated combinations of constructs were cotransformed into yeast cells (strain AH109) and grown on the nonselective medium SD/-L-W and selective medium SD/-L-W-H.

**B.** A split complementary luciferase assay confirmed the interactions between OsNF-YB7 and OsGLK2. Coexpression of the fusion OsGLK2 and the N-terminal half of LUC (nLUC-GLK2) and the fusion of C-terminal half of LUC and OsNF-YB7 (cLUC-NF-YB7) in the epidermal cells of *N. benthamiana* leaves induced LUC activities, whereas the epidermal cells coexpressed nLUC-GLK2 and cLUC, nLUC and cLUC-NF-YB7, or nLUC and cLUC did not show LUC activities.

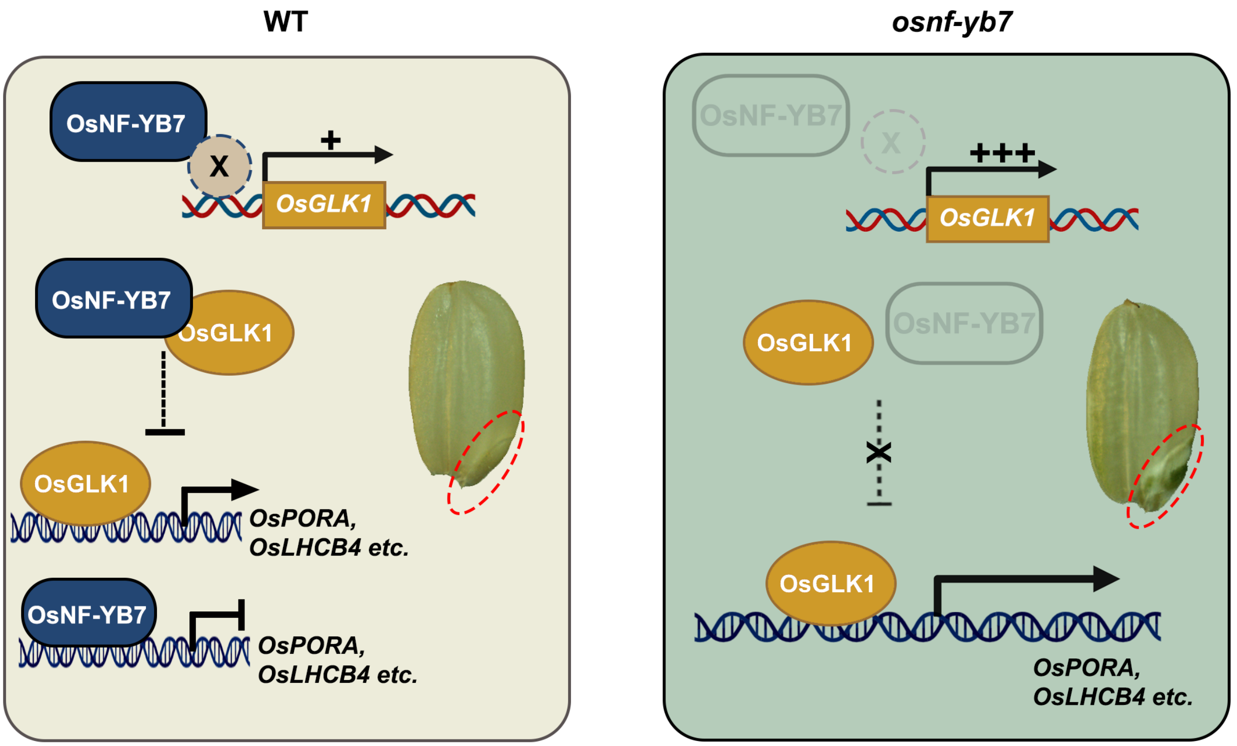


**Supplementary Figure 14. A proposed model of the OsNF-YB7-mediated suppression of Chl biosynthesis in rice embryo.**

In WT, OsNF-YB7 transcriptionally represses *OsGLK1*, probably requiring the assistance of an unidentified transcription factor X to recognize the promoter of *OsGLK1*. On the other hand, OsNF-YB7 can interact with OsGLK1 to repress its transactivation activity on downstream photosynthesis and Chl biogenesis genes like *OsPORA* and *OsLHCB4*. In addition, OsNF-YB7 can also directly bind to the promoters of *OsPORA* and *OsLHCB4*, consequently repressing their transcription. In the *osnf-yb7* mutant, OsGLK1 is activated to promote the expression of downstream Chl biosynthesis- and photosynthesis-releated genes, consequently inducing chloroembryo.
